# Automated Discrimination of Dicentric and Monocentric Chromosomes by Machine Learning-based Image Processing

**DOI:** 10.1101/037309

**Authors:** Yanxin Li, Joan H. Knoll, Ruth Wilkins, Farrah N. Flegal, Peter K. Rogan

## Abstract

Dose from radiation exposure can be estimated from dicentric chromosome (DC) frequencies in metaphase cells of peripheral blood lymphocytes. We automated DC detection by extracting features in Giemsa-stained metaphase chromosome images and classifying objects by machine learning (ML). DC detection involves i) intensity thresholded segmentation of metaphase objects, ii) chromosome separation by watershed transformation and elimination of inseparable chromosome clusters, fragments and staining debris using a morphological decision tree filter, iii) determination of chromosome width and centreline, iv) derivation of centromere candidates and v) distinction of DCs from monocentric chromosomes (MC) by ML. Centromere candidates are inferred from 14 image features input to a Support Vector Machine (SVM). 16 features derived from these candidates are then supplied to a Boosting classifier and a second SVM which determines whether a chromosome is either a DC or MC. The SVM was trained with 292 DCs and 3135 MCs, and then tested with cells exposed to either low (1 Gy) or high (2-4 Gy) radiation dose. Results were then compared with those of 3 experts. True positive rates (TPR) and positive predictive values (PPV) were determined for the tuning parameter, σ. At larger σ, PPV decreases and TPR increases. At high dose, for σ= 1.3, TPR = 0.52 and PPV = 0.83, while at σ= 1.6, the TPR = 0.65 and PPV = 0.72. At low dose and σ = 1.3, TPR = 0.67 and PPV = 0.26. The algorithm differentiates DCs from MCs, overlapped chromosomes and other objects with acceptable accuracy over a wide range of radiation exposures.

## Introduction

Clastogenic events producing dicentric chromosomes (DC) are among the most reliable biomarkers of radiation exposure. These events are infrequent relative to the background of normal monocentric chromosomes (MC), thereby requiring many cells for accurate dose estimation. This has motivated efforts to automate cytogenetic image analysis. This task has been a longstanding challenge in computer vision research (Bayley et. al. 1991), largely because chromosome morphology is incredibly variable between metaphase cells and different preparations and laboratories. The reasons include differences in chromosome structure, staining methods, biological effects and differences in sample preparation methods. Metaphase cell selection strongly influences the accuracy of these analyses. Content and classification-based methods have been used to rank metaphase cell images based on chromosome number and degree of chromosome overlap (Kobayashi et al. 2004). Nevertheless, advances in automated karyotyping have been limited by the accuracy of algorithms, and hidden implementation details of commercial products.

Spurious branches produced by medial axis thinning of irregular chromosome objects can lead to incorrect centromere placement. We developed an algorithm to calculate the centerline of the chromosome that excluded spurious branches and was independent of overall morphological differences (Subasinghe et al. 2010; Subasinghe et al. 2013). This approach spurred new strategies for centromere detection using curvature rather than width to determine centromere location (Mohammaed 2012) or artificially straightened chromosomes to create a trellis perpendicular to the centerline (Jahani and Setarehdan 2012). However, these methods, including our own, require objects with smooth chromosomal boundaries. The presence of irregular contours adversely impacts the centreline, and consequently, the accuracy of features used to infer centromere location. Centerline-based results are also affected by chromosomes exhibiting sister chromatid separation (SCS).

Metaphase images containing ∽46 individual, non-overlapped chromosomes without SCS will yield the most accurate DC detection. In practice, such ideal images are uncommon among cell preparations in biodosimetry laboratories so a method of selecting appropriate metaphases or dealing with overlaps is required. In this manuscript, we present a series of image processing methods to automate detection of DCs. The process involves selecting metaphase cells with optimally distributed chromosomes (Kobayashi et al. 2004) from a sample, defining the boundaries of the remaining chromosomes, detecting centromere candidates, and discriminating mono-from dicentric chromosomes. When multiple chromosomes overlap or touch in an image, these clusters are preprocessed and separated by a watershed transform, which ensures that valid chromosome objects are processed.

The method segments the chromosome objects using local thresholding and draws object outlines by Gradient Vector Flow (GVF) active contours (Xu and Prince 1998). Once the object is extracted based on the GVF outline, the contour of the chromosome is partitioned along the centreline using a polygonal shape simplification algorithm called Discrete Curve Evolution (DCE) (Latecki and Lakamper 1999, Bai et al. 2007).

We recently implemented a centromere localization algorithm, which is refractory to the confounding effects of highly bent chromosomes and SCS (Subasinghe et al.2015). Since centerline-based centromere detection tends to perform better than other approaches, the centerline is used to partition the chromosome contour into two nearly symmetric regions. The centerline is not used to measure chromosome width or other properties. As a result, the boundary texture does not affect the smoothness of the width profile measurements which are used to locate centromeric constriction(s). Once the contour is partitioned and segmented, an Intensity Integrated Laplacian (IIL) thickness measurement algorithm (Subasinghe et al. 2013) integrates pixel intensities, resulting in vectors traced axially along homogenous intensity regions, analogous to chromosome bands. Here, we derive features in chromosome images to rank centromere candidates by Support Vector Machine (SVM) learning. These features represent various aspects of the geometry and other properties of the chromosome at the locations of the selected candidates. A second SVM is then used to discriminate monocentric and dicentric chromosomes.

## Materials and Methods

The algorithm and software separates and isolates chromosomes, localizes centromere candidates within each, then processes the candidates to distinguish MCs from DCs. This is done by extracting valid chromosomes from images of complete metaphase cells using customized image-processing methods, computing quantitative features from these images as input to pretrained ML models that optimize identification of DCs among a larger population of MCs.

### Image Segmentation

All objects in images are first segmented and binarized by local intensity thresholding (Otsu 1979). The foreground objects obtained are a mixture of single chromosomes, clusters of overlapped or touching chromosomes, nuclei and staining debris. Touching and overlapped chromosome clusters are problematic for DC analysis as their inclusion presents multiple centromeres in one object. To separate chromosome clusters into individual chromosomes, we perform a watershed-based method. The watershed transform, a widely used technique in image segmentation (Meyer 1994), treats an image as a surface and consequently finds catchment basins and ridge lines that separates domains of the object. The transform is guided by seeds placed by users to match possible basins on the image. Aggressive intensity re-thresholding on foreground pixels is calculated for all objects. New segmented regions act as seeds in the watershed transform. Therefore, the ridge pattern combines intensity and positioning information, which provides a possible separation strategy for the object (Figure 1A). However, single chromosomes with considerable SCS or non-uniform staining can also be broken at the site of a ridge pattern. Fragments caused by incorrect splitting exhibit different morphological characteristics from complete chromosomes. We established three simple empirical conditions based on feature length, perimeter and area to prevent inappropriate splitting of chromosomes (Figure 1B). Ridges that meet any of the conditions are considered to split a single chromosome and are therefore discarded. The two parts of an object separated by a ridge (R) are referred to as *P* and *Q*.

**Figure 1.**
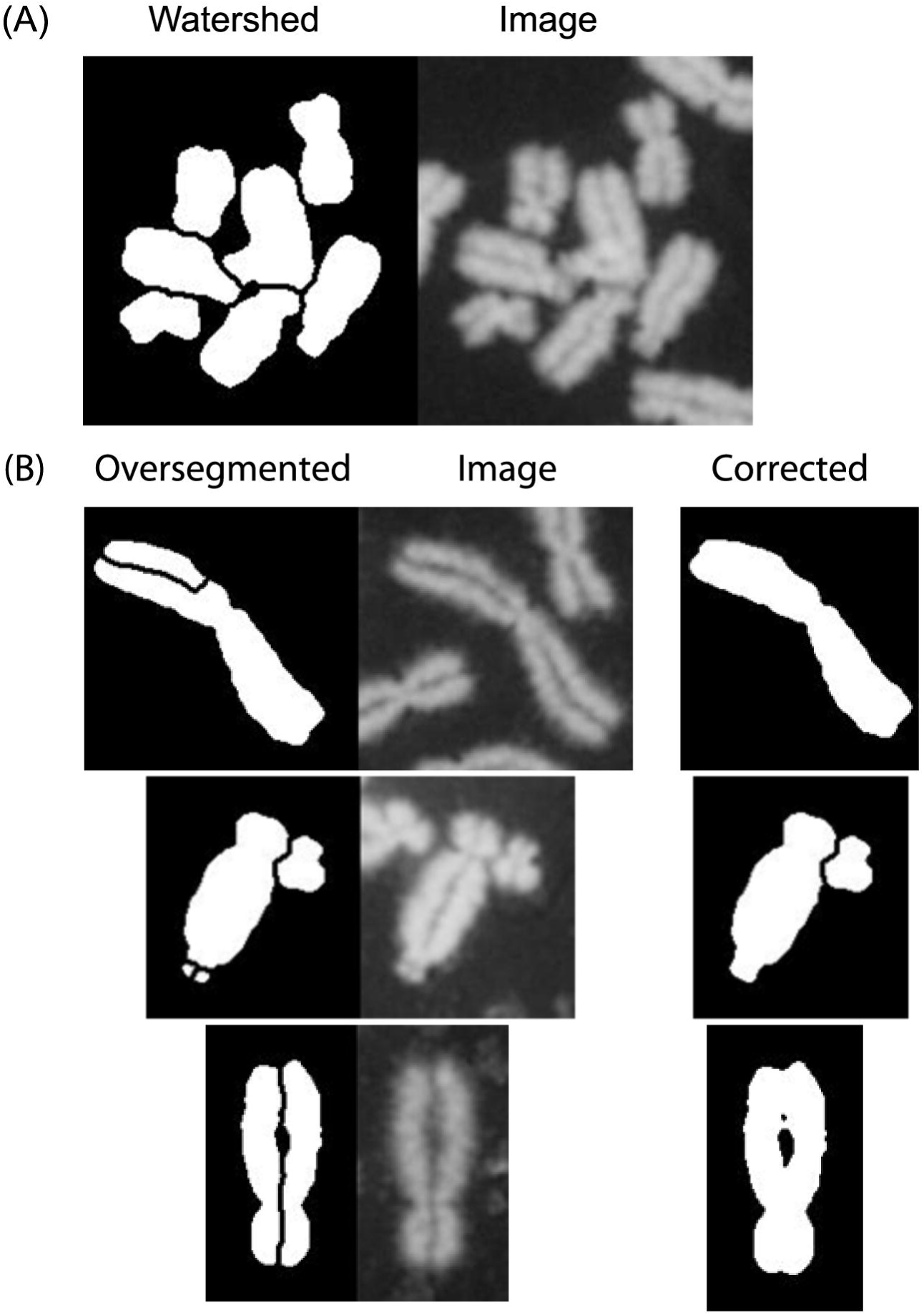
Modified watershed separation of chromosome clusters. After the original metaphase image is binarized by intensity threshold Segmentation, connected chromosome clusters are formed due to under-segmentation. Panel (A) Watershed separation operation is applied to these clusters to prevent oversegmentation. This involves determining the lengths of the ridges between components of the cluster, areas of the separated regions, and the degree of symmetry of the separated regions; (B) Constraints are applied to prevent oversegmentation of individual chromosomes if: i) length of the ridge exceeds half of the perimeter of one of the separated regions, ii) areas of small regions separated by the operation are less than 10% area of the larger region, and iii) The two separated regions exhibit highly symmetric structures adjacent to the ridge the separates them.

Condition 1: *R_length_* > 0.55 * *min* {*P_perimeter_, Q_perimeter_*}.
Condition 2: *min*{*P_area_, Q_area_*} < 0.1 * *min*{*P_area_, Q_area_*}.
Condition 3: 85% of *P, Q*’s area are spatially symmetric with *R* being the axis and R_length_ > 0.3 * min {*P_perimeter_, Q_perimeter_*}.

Conditions 1 and 2 are designed to avoid breaking of complete chromosomes. Condition 3 prevents splitting of sister chromatids. All parameters for these conditions have been heuristically chosen and validated with large numbers of images containing touching and overlapping chromosomes. However, separation of these objects cannot be guaranteed.

To filter out non-chromosomal objects, we examined the sizes, brightness and contours after segmentation of all objects in an image. Upper and lower thresholds for chromosome area and average intensity have been determined from statistical distributions of these values from analysis actual chromosomes in a set of metaphase cells. Chromosome fragments, nuclei and staining debris are eliminated if they are respectively above or below the thresholds for either chromosome area (>5x the area of neighboring median object size or <200 pixels^2^) or intensity (>20x mean intensity of median size objects or <40x mean intensity of median size objects). To detect overlapping chromosomes and other unfiltered chromosomal objects in the image, the contour of each object was analyzed. We measure the point-wise inner distances (Ling and Jacobs 2007) of the contour to estimate the maximum width of a chromosome. DCE simplified contours are used, replacing original contours to reduce computational time complexity. Outliers of the estimated width in a metaphase are removed as overlapped chromosomes.

### Centromere Localization

Chromosomes are serially processed by the GVF, DCE and the IIL algorithms [Subasinghe et al. 2013], then candidate centromeres are selected from local minima along the width profile of each chromosome. A Support Vector Machine (SVM) was previously trained on 11 image analysis features (Subasinghe et al. 2015) to find the strongest candidate centromere with the based on its distance to the hyperplane relative to all others. Briefly, these features describe: i) the local minimal chromosome width, the pixel intensity at each candidate; ii) differences between a curve fit to the width profile and the profile itself; iii) the maximal width adjacent to the candidate; iv) the beginning and end coordinates of the Intensity Integrated Laplacian vectors, v) the shortest distance from the candidate to the end of the centerline; and vi) the ratio of width at the candidate to the average width of all points along the centerline.

This centromere SVM identifies a single candidate as the centromere, regardless of whether the chromosome is MC or DC. To identify secondary centromere candidates, the top candidates are sorted in order of their signed distances to the SVM hyperplane and the two best candidates are then analyzed. The true centromere(s) are expected to be present among the candidates. In the case of a MC chromosome, the two candidates comprise a true centromere and a non-centromeric region; for DC chromosomes, both candidates would include the true centromeres. To improve the accuracy of centromere assignment, it was necessary to incorporate 3 additional image features (A1 – A3, defined below) in the centromere SVM, defined as follows. For each chromosome, let *c_i_*, 1 ≤ *i* ≤ *N* denote the *i_th_* point on its centerline. We introduce the following notations:

*I(c_i_)* refers to the image intensity value at *c_i_*.
*W(c_i_)* and *W(c_i_, c_j_)* refer to the width profile at *c_i_*, or of the interval [*c_i_, c_j_*].
*W′(c_i_)* refer to the quadratic curve fit to the width profile at *c_i_*.
*l_s_ (c_i_)* and *l_e_ (c_i_)* refer to Laplacian start and end points corresponding to *c_i_*.

For each centromere candidate *k* of the same chromosome, *c_k_*, the additional features are described below:

A1: *I(c_k_)/MAX(I(c_i_), i* = 1, 2, 3… *N*). This is the normalized intensity of the candidate.
A2: ∠(*l_s_(c_k_), c_k_, l_e_(c_k_)*). This feature is the turning angle between the start and endpoints of the Intensity Integrated Laplacian vector at the candidate. A3: *W′(c_k_)* – *W(c_k_)*. The difference of the fitted quadratic width and the actual width of the candidate.

Feature A1 extracts intensity values at the centromere candidates. Feature A2 prevents false candidates at bending or twisting regions in a chromosome. The width profile of a chromosome contains a set of discrete width values with peaks in the middle and valleys at the ends of each which are fit to a quadratic function. Centromeres normally show significant reduction in width due to constrictions at these contour coordinates. This chromosome property can be captured by comparing the actual width profiles at the centromere candidates to their expected widths fit to the quadratic function. Feature A3 in the centromere SVM measures the difference between these values. Along with the original features, the final centromere SVM uses 14 features to select the optimal candidates used in the detection of DCs.

### DC Detection

A compound ML model was developed to discriminate MCs from DCs. The components of the model included a second SVM trained to recognize MCs and DCs, whose accuracy was enhanced with a Boosting Classifier (Viola and Jones 2001). Given the two candidate centromeres, the method generates a set of features for a chromosome which characterize their respective impacts on chromosome structure. We developed a set of image features (F1 – F16, defined below) to train the MC-DC SVM to distinguish between them. In a DC, each candidate is expected to exhibit a constriction of similar magnitude, but their respective widths will differ in MC chromosomes. The MC-DC SVM analyzes selected candidates in the context of the chromosome. Significant variation between the morphologies of different chromosomes required some features to be designed to mitigate the occurrence of false positive DCs, which were, in fact, true MCs. To illustrate these features, we use *c_i_*, 1 ≤ *i* ≤ *N* to denote the *i_th_* point along the centerline of a chromosome. In addition to the expressions defined above, we also introduce the following symbols:

*E(c_i_, c_y_)* refers to the normalized accumulated Euclidean distance between *c_i_* and *c_j_* along the centerline.
*H(C_i_)* refers to the distance from *c_i_* to the hyperplane in the centromere SVM, if it is a candidate.
*D_s_(C_i_)* and *D_e_(c_i_)* refer to *c_i_*’s Euclidean distances to *l_s_(c_i_)* and *l_e_(c_i_)*.
μ and σ denote the mean and standard deviation, respectively, for sample distributions.

We define the selected centromere candidates as *c_p_* and *c_q_*, with *p* < *q*, and summarize features based on these candidates in the MC-DC SVM below:

F1, F2: *H(c_p_)* and *H(c_q_)*. They are the likelihoods of the candidates being true centromeres evaluated by the centromere SVM.
F3: | *H(c_p_) – H (c_q_)* |. DC chromosomes should have similar F1 and F2 values since both candidates are true centromeres and a smaller F3 value. By contrast, in MC chromosomes, F3 tends to be large, as one of the candidates is a false centromere.
F4: *E(c_p_, c_q_)*. This feature prevents cases where the two candidates are so close that they actually belong to the same centromere.
F5: *min* {*E*(*c_i_, c*_1_), *E*(*c_i_, c_N_*), *E*(*c_j_, c*_1_), *E*(*c_j_, c_N_)*}. This feature prevents false positive cases where a candidate is positioned too close to telomeres.
F6: *W_max_* (*c*_1_, *c*_*p*-1_) + *W_max_* (*c*_*p* + 1_, *c_N_*) – 2 × *W*(*c_p_*). This feature is part of the centromere SVM.
F7: *W_max_* (*c*_1_, *c*_*q* - 1_) + *W_max_* (*c*_*q*+1_, *c_N_*) – 2 × *W*(*c_q_*). F6 and F7 measure the contour constriction at the centromere candidates.
F8: *max*{*Z*(*c_p_*), *Z*(*c_q_*)}, *Z*(*x*) = (*W*(*x*) – *W_μ_*(*c_p_, c_q_*))/*W_σ_* (*c_p_, c_q_*). This feature is the larger value of the z-scores for the candidates’ width profiles. It is relatively small for DC chromosomes, and large for MC chromosomes.
F9: *min* {*A, B*}/*max*{*A, B*}, where *A* = *W*(*c_p_*) – *W_μ_*(*c*_*p*+1_, *c*_p+3_), *B* = *W*(*c_q_*) – *W_μ_* (*c*_*q* − 3_, *c*_*q* − 1_). This feature assesses the similarity of the steepness at the candidate locations on the chromosome.
F10: |*R*(*c_p_*) – *R*(*c_q_*)|, *R*(*x*) = *min* {*D_s_*(*x*), *D_e_* (*x*)}/*max* {*D_s_* (*x*), *D_e_* (*x*)}. This feature detects false centromeres that are caused by chromosome bending.
F11, F12: *θ_s_*(*p*) and *θ_s_*(*q*), where *θ_s_*(*x*) = ∠ (*l_s_*(*c*_*x*−5_), *l_s_*(*c_x_*), *l_s_*(*c*_*x*+5_)). These features detect the contour concavities of the Laplacian start points for the candidates.
F13, F14: *θ_e_*(*p*) and *θ_e_*(*q*), where *θ_e_*(*x*) = ∠ (*l_e_*(*c*_*x*−5_), *l_e_*(*c_x_*), *l_e_*(*c*_*x*+5_)). These features detect the contour concavities of the Laplacian end points for the candidates.

Features derived from width profiles and contours are founded on the knowledge of cytogenetic characteristics of centromeres, which are specifically associated with the analysis of DCs.

However, the diversity of raw intensity pixel values between different chromosomes and images discourages the use of unprocessed features in these supervised learning models. This problem was addressed with generalized pixel-level features that are widely used in various recognition-driven problems in computer vision. A Boosting Classifier applied to Haar-like features in chromosome images uses this pixel-level information to strengthen the accuracy of centromere probability measurement (Viola and Jones 2001). Haar-like features have been proven to be an effective descriptor for low-level intensity patterns. Pixel intensities are integrated in moving sub-windows and the integrated values are compared within windows comprising a series of symmetric rectangles. This mechanism generates a comprehensive gray-scale descriptor for a region of interest. In most applications, Haar-like features work with Boosting classifiers because of the high dimension of the feature set. A Boosting model consists of a large number of simple classifiers that are only required to be more accurate than a random classifier. During training, the Boosting model iteratively adjusts weights of its classifiers, and combines all classifier predictions to improve accuracy. The sign of the weighted sum of the Boosting classifiers determines the binary classification. Haar-like features, computed in a 21-by-21 region centered on a selected candidate, comprise 6749 features input to the Boosting classifier. The weighted sums of the classifier of both candidates in a chromosome are appended to the MC-DC SVM as additional features (F15, F16). Various Boosting configurations (eg. Ada Boost and Robust Boost) were also tested to determine if these improved discrimination of candidate centromeres.

The performance of different kernel types, linear, polynomial and radial basis function (RBF) kernels, were compared for the MC-DC SVM. The centromere SVM was previously configured to use the RBF kernel [Subasinghe et al. 2015]. Similarly, RBF was selected for the MC-DC SVM classifier, due to its superior accuracy in distinguishing MCs and DCs in a curated set of chromosomes (see Results). SVMs can produce multiple classifier models, each based on a unique tuning parameter, σ. Increasing σ values effectively represent a tradeoff between increased sensitivity and reduced specificity in DC detection. The RBF is tuned with the parameter, whose value monotonically increases (1.1 – 1.8) with increased detection of DCs (both true and false positives [TP, FP]). The optimal results are determined by testing these values. For example, the inferred DC distribution in a sample at different values of is fit to the expected Poisson distribution of DCs in irradiated lymphocytes [International Atomic Energy Agency 2001].

### Software Organization

The algorithms were originally developed in MATLAB, and the finished software has been implemented in C++. The current version has been re-organized from its last release, logically divided into four layers. The architecture is indicated in Figure 2.

**Figure 2.**
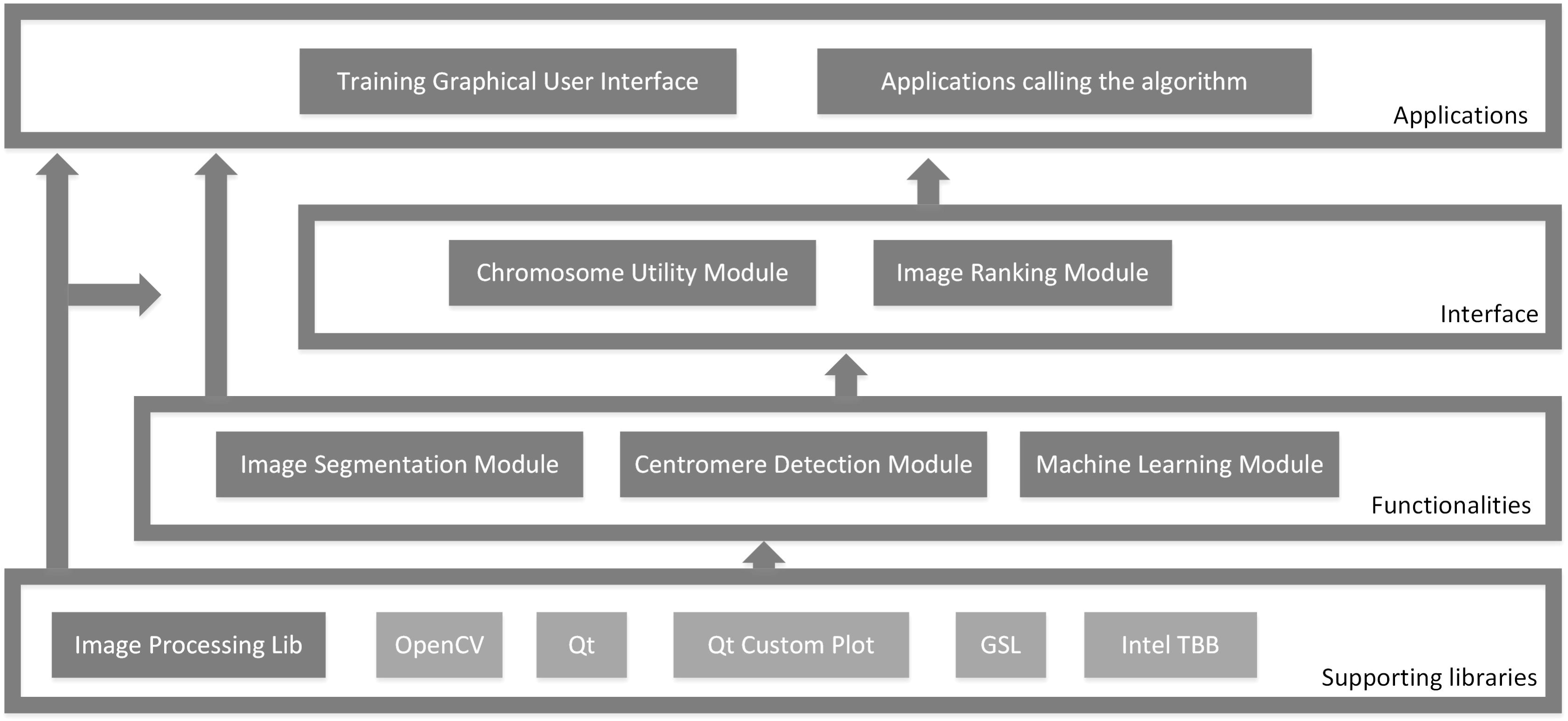
UML diagram of software development system. The figure illustrates the structure of the chromosome image processing software system in a layered structure based on functional modules called during training and testing of the SVM components. Software modules are the displayed as boxes and containing rectangles represent development layers. Light gray boxes and dark gray boxes indicate third-party libraries and libraries developed by our team respectively. Software building dependencies are showed by arrows. The layers supporting libraries, functionalities and interface comprise the complete automated dicentric chromosome identification algorithm. Any application using the algorithm belongs to the applications layer, including our training graphical user interface.

The supporting libraries layer includes third-party libraries, as well as low-level image processing modules. Most core classes and functions are built on OpenCV and Qt libraries. Intel Thread Building Blocks (TBB) provides multi-threading parallel processing for DC analysis operations. The GNU scientific (GSL) and Qt libraries are also called by the software. The main DC analysis is implemented in the functionalities layer and contains three modules corresponding to the three stages of the ADCI algorithm: image segmentation, centromere detection and ML. We create the interface layer as an intermediate between DC analysis and user interfaces. Core data structures and classes representing metaphase images, chromosomes and other key cytogenetic concepts are coded in this layer.

The top tier is the applications layer, including multiple applications depending on the end user requirements. A graphical user interface (GUI) was developed to obtain training data for the SVMs. This GUI supports user scoring by visually displaying the centromere candidates on each chromosome. These data are compared with ground truth-scored centromeres by the training GUI to assess performance of the SVM iterations and feature improvements during the development process. A version of this software application can be used to evaluate individual DC and MC chromosomes either in the available image gallery or supplied by the user (Figure 3).

**Figure 3.**
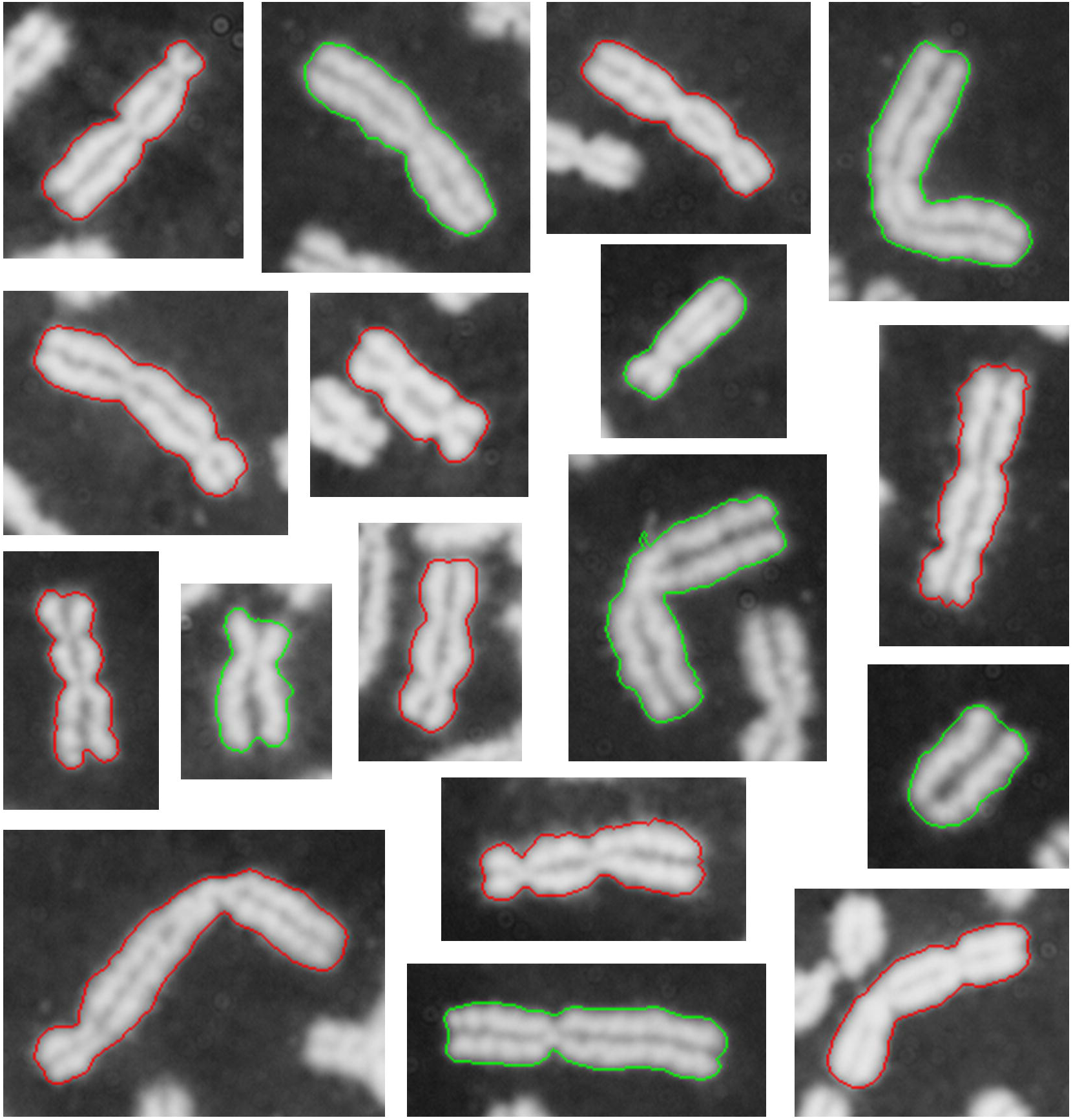
Classification of mono- and dicentric chromosomes. The figure displays a representative set of MCs and DCs, as well as the classification results scored by the MC-DC SVM (sigma=1.5). The contour of the chromosome defined by the algorithm is color coded as either monocentric (green) or dicentric (red). Chromosomes are cropped from metaphase images in a sample exposed to a 3-Gy X-ray radiation source provided by CNL.. These examples can be classified with the centromere and MC-DC SVMs online with a software application available at http://cytobiodose.cytognomix.com.

## Results

### Data sources

Unlike the centromere detection procedure, most experimental data analyzed are from cells that have been exposed to calibrated gamma or X-ray radiation sources. The microscopy images of metaphase cells were generated in biodosimetry laboratories at Health Canada (HC) and Canadian Nuclear Laboratories (CNL). Experts in these laboratories determine the biological level of radiation exposure in accidents and other exercises. The datasets were comprised of multiple batches of images from samples of different known radiation exposures (from 1-4 Gy). Cytogenetic experts collected chromosome information for routine biodosimetry exercises, which have been used to develop and test the automated methods described in this study. Distinct datasets were used to derive the ML models and to evaluate their performance by crossvalidation. An early version of the software was used to record key attributes used for training, ie. 3 experts marked all true centromeres amongst the set of candidates on each DC chromosome, and denoted false positive DCs.

Cytogenetic specialists at UWO, HC, and CNL used the graphical user interface version of the software (Figure 2), which provided training data for the SVM that indicated ground truth designations of dicentric, and in some instances, monocentric chromosomes. Chromosomes were first classified by a SVM; then, users scored chromosomes as DC or MC by confirming or correcting this classification. Scoring differences resulted from SVMs with different sigma values (1.4 vs. 1.5), and scoring criteria adopted by different specialists. For example, the classification of dicentric acrocentric chromosomes depends on the length of the p arm and the proximity of. the centromere to the nearest telomere. If this distance is particularly short, i.e. less than the chromatid width, a potential DC is not counted as dicentric, as the determination is ambiguous for the software. Differences between scores were then discussed and usually could be resolved by joint review. Any remaining discrepancies are reported in the final results.

The metaphase image data were divided into 3 groups, according to how each was scored. Cytogenetic experts scored all DCs in each dataset. Dataset 1 contained 281 fully labeled metaphase images with centromeres marked by experts. 266 DC chromosomes, 3,222 MC chromosomes are present in dataset 1, with all other segmented objects being chromosome clusters, nuclei and staining debris. In dataset 2, only true DC chromosomes are scored while other objects, including MC chromosomes, are ignored. In dataset 2, we observed 531 DC chromosomes and 13,898 other objects from 612 images. Both datasets 1 and 2 are from cells exposed to 3-4 Gy (high-level) gamma radiation. The image segmentation of these datasets was subjected to intensity thresholding without application of the watershed method. The final dataset 3, comprises a wide range of doses and has been separated into 1 Gy (low dose) and 3-4 Gy high dose subsets. This dataset 3 was analyzed with a version of the algorithm that included watershed segmentation.

### Image Segmentation

The watershed separation and the segmentation components were tested with an dataset enriched in chromosome clusters created from 60 metaphase images from dataset 1. It contained 2340 objects including 1762 single chromosomes, 349 chromosome clusters and 229 nuclei and debris or fragmentary objects. The watershed method separates 294 chromosome clusters, or 84% of the set of 349. Some single chromosomes (n=48) were inappropriately broken by the watershed method, however 1714 (97%) remained intact. A portion of whole nuclei, fragments and debris objects (n=84) were also split by the watershed method, however none of these were classified in subsequent steps as either MC or DCs.

### Centromere SVM

The centromere SVM model in our DC analysis selected centromere candidates to provide information to assign the type of chromosome by the MC-DC SVM. We evaluated the performance of the centromere SVM on the basis of selected candidates that identified true centromeres. Only DCs were assessed, as it was very rare that the centromere in a MC was not among the two candidates. The detection accuracy based on the 2 most highly-ranked centromere candidates in a chromosome was compared with the 4 top-ranked candidates. Both centromeres in a DC were required to be identified in either the top 2 or top 4 candidates. In dataset 1, a 5fold cross-validation was carried out with 4 of 5 DCs defined as training data and the remainder were used for testing the SVM Subsequently, the full centromere SVM was trained with all DCs in dataset 1, and tested with data from dataset 2 (results are shown in Table 1).

**Table 1.** Performance of centromere SVM

|  | Cross-validation in dataset 1 | Testing of dataset 2 |
| --- | --- | --- |
| Total no. of DCs present | 266 | 531 |
| No. DCs detected with top 2 candidate centromeres (%) | 194 (73%) | 371 (70%) |
| No. DCs detected with top 4 candidate centromeres (%) | 248 (93%) | 499 (94%) |

### Boosting and the MC-DC SVM

We applied several types of Boosting classifiers, which combine different features to improve the performance of weak SVMs. We compared the performance of Boosting models available in the MATLAB Image Processing Toolkit and the C++ OpenCV library. Boosting classifiers were trained using selected candidates of chromosomes in dataset 1, including 6906 candidates comprising both DC and MC chromosomes. The Boosting models were assessed by comparing results from Adaptive Boosting in OpenCV, as well as Adaptive Boosting and Robust Boosting in MATLAB. The lowest accuracy, 87%, was found using Adaptive Boosting method in MATLAB, whereas the Adaptive Boosting in OpenCV exhibited a slightly higher accuracy (89%). The results demonstrate that various Boosting models have highly similar training accuracies and therefore, we do not discriminate between them.

For the MC-DC SVM, we evaluate combinations of candidate centromeres produced by the centromere SVM for individual chromosomes. The number of TP DCs and the number of MCs incorrectly labeled positive (FPs) by the SVM are assessed by expert review. The PPV (also called precision) and TPR (also known as sensitivity or recall) are used to assess the performance of the SVM at different σ values. PPV indicates the exactness of DC detection. TPR measures the fraction of true DC detection. We seek feature sets and σ values that maximize PPV and TPR using the same training data. Since the MC-DC SVM is limited by the selections made by the preceding centromere SVM, the centromere SVM trained with the complete dataset 1 is used to provide selected candidates. Only DC chromosomes with both centromeres selected are counted towards correct proportion of DCs classified.

The model derived from dataset 1 was evaluated by cross-validation. The centromere SVM made correct selections for 194 of the 266 DCs. A Boosting classifier was trained by 5 fold crossvalidation, followed by sequential training of the MC-DC SVM with the same cross-validation schema. The Boosting-SVM model was then tested. Results shown in Table 2 indicate that the σ value of 1.4 achieves the highest combined PPV and TPR.

**Table 2.** Results of MC-DC SVM cross-validation on dataset 1

| Sigma | TPs | FPs | PPV% | TPR<sup>1</sup> % | TPR<sup>2</sup> |
| --- | --- | --- | --- | --- | --- |
| 1.0 | 91 | 18 | 83.5 | 46.9 | 34.2 |
| 1.1 | 111 | 24 | 82.2 | 57.2 | 41.7 |
| 1.2 | 124 | 28 | 81.6 | 63.9 | 46.6 |
| 1.3 | 134 | 35 | 79.3 | 69.0 | 50.4 |
| 1.4 | 148 | 41 | 78.3 | 76.3 | 55.6 |
| 1.5 | 154 | 49 | 75.9 | 79.4 | 57.9 |
| 2.0 | 166 | 79 | 67.8 | 85.6 | 62.4 |

<sup>1</sup> Total of 371 chromosomes with both centromeres correctly detected by Centromere SVM;
<sup>2</sup> Total of 531 chromosomes with all known DCs scored, regardless of results of Centromere SVM

In addition to cross-validation, we also tested dataset 2 using a Boosting-SVM model that was trained using dataset 1. By contrast with dataset 1, MC chromosomes were not scored or labeled in dataset 2. Since MC-DC SVM distinguish DC from non-DC objects, and the non-DC objects comprise a mixture of MCs, intact nuclei, debris and acentric fragments, this is actually a more stringent evaluation than the original approach. The centromere and MC-DC SVMs correctly selected 371 of the 531 DCs present (Table 3).

**Table 3.**
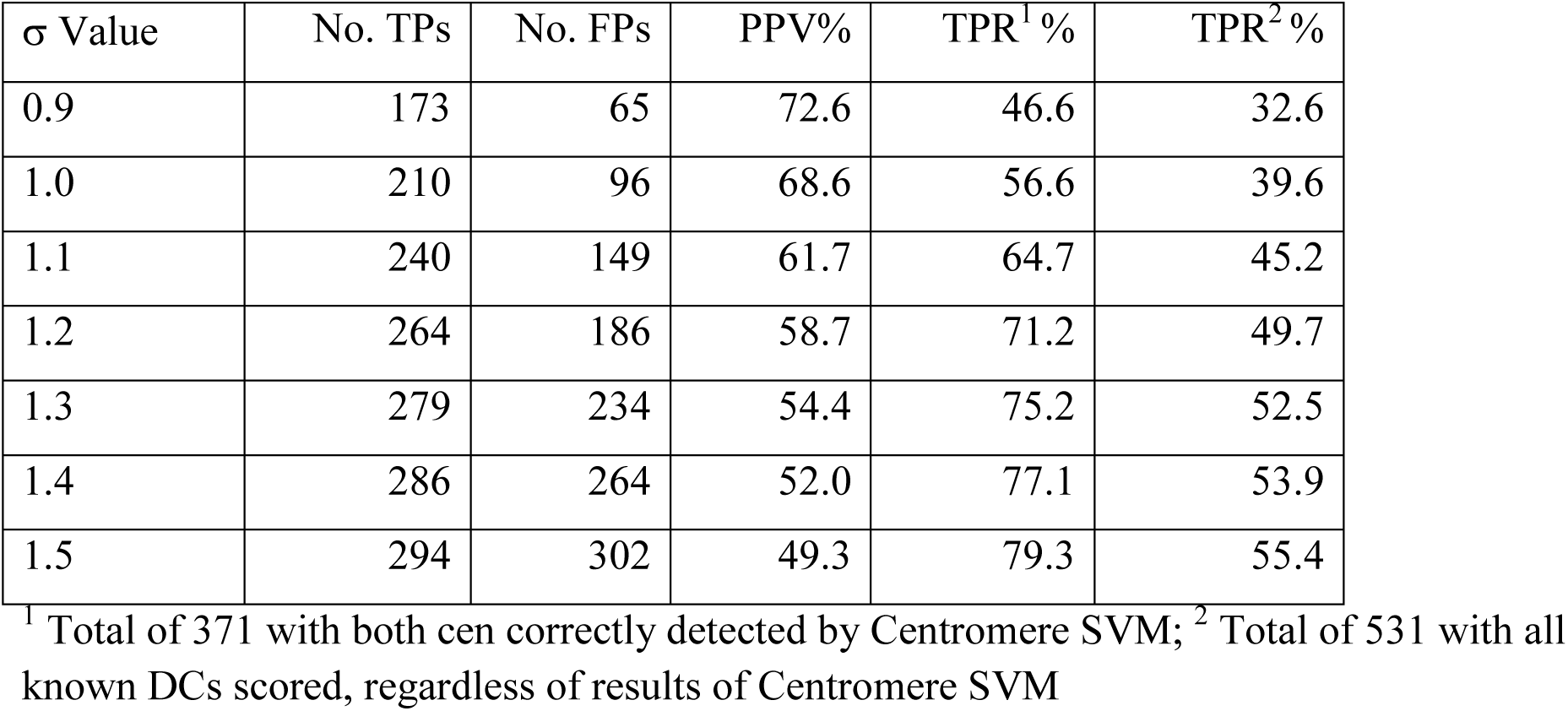
Results of MC-DC SVM test on dataset 2

| $\sigma$ Value | No. TPs | No. FPs | PPV% | TPR<sup>1</sup>% | TPR<sup>2</sup>% |
| --- | --- | --- | --- | --- | --- |
| 0.9 | 173 | 65 | 72.6 | 46.6 | 32.6 |
| 1.0 | 210 | 96 | 68.6 | 56.6 | 39.6 |
| 1.1 | 240 | 149 | 61.7 | 64.7 | 45.2 |
| 1.2 | 264 | 186 | 58.7 | 71.2 | 49.7 |
| 1.3 | 279 | 234 | 54.4 | 75.2 | 52.5 |
| 1.4 | 286 | 264 | 52.0 | 77.1 | 53.9 |
| 1.5 | 294 | 302 | 49.3 | 79.3 | 55.4 |

<sup>1</sup> Total of 371 with both cen correctly detected by Centromere SVM; <sup>2</sup> Total of 531 with all known DCs scored, regardless of results of Centromere SVM

Dicentric chromosomes (FNs) missed in dataset 2 were then reclassified and appended to the DC training data as TPs, the MC-DC SVM was retrained, and then tested on independent dataset 3. A cytogenetic expert in our research group (JHMK) scored DCs of all metaphase cells in dataset 3 as ground truth. Specialists from HC and CNL also scored a common subset of 144 of these metaphases in the high-dose subset for comparative study. Comparison of the retrained model with the ground truth scoring indicated retraining the model significantly increased the PPV (approximately 20%).

In the high dose exposure subset, the software segmented 14428 objects, averaging 40 objects per metaphase. Our UWO expert (JHMK) designated 476 objects as DCs, with 179 in the 144 metaphase cells scored by all experts. At low-dose (1 Gy), the software detected 8,041 objects, an average of 38.7 objects per image. The DC chromosomes in cells exposed to low dose radiation are infrequent. The expert (JHMK) found 27 DC chromosomes in the low-dose subset. The comparison of the MC-DC SVM with ground truth and inter-specialist comparisons are shown in Table 4. The results are stratified according to (a) a subset of DCs from cells exposed to high dose radiation scored by all experts and compared those produced by the software, (b) all high dose DCs identified by the software relative to scoring by JHMK, and (c) DCs detected in a low dose sample compared to JHMK’s interpretation. Using σ of 1.4 or 1.5, at high dose exposures, approximately half of DCs are detected with acceptable false positive rates (PPV = 71 - 77%). At low dose in which fewer DCs form, sensitivity of detection is higher (66-74%), at a cost of significantly lower specificity (PPV = 18 – 21%), the latter being related to quality of the data and current limitations of the algorithm. Scoring of DCs of different experts were minimally discordant (<3%).

**Table 4.** Performance of MC-DC SVM on dataset 3, consisting of metaphase cells subject to high-dose and low-dose exposure: comparison with expert scoring by University of Western Ontario (UWO), Health Canada (HC) and Canadian Nuclear Laboratories (CNL)

| Source of dicentric chromosome data | No. chromosomes; Percentage | SVM $\sigma$ value | | | | | | | HC | CNL | UWO<sup>1</sup> |
| --- | --- | --- | --- | --- | --- | --- | --- | --- | --- | --- | --- |
|  |  | 1.2 | 1.3 | 1.4 | 1.5 | 1.6 | 1.7 | 1.8 |  |  |  |
| High-Dose chromosome data, commonly scored<sup>2</sup> | TPs | 71 | 79 | 90 | 98 | 102 | 108 | 110 | 175 | 176 | 179 |
|  | FPS | 13 | 17 | 33 | 39 | 46 | 54 | 66 | 4 | 3 | 0 |
|  | PPV% | 84.5 | 82.3 | 73.2 | 71.5 | 68.9 | 66.7 | 62.5 | 97.8 | 98.3 | 100 |
|  | TPR% | 39.7 | 44.1 | 50.3 | 54.8 | 57.0 | 60.3 | 61.5 | 97.8 | 98.3 | 100 |
| All High-Dose chromosome data<sup>3</sup> | TPs | 214 | 250 | 280 | 301 | 314 | 327 | 333 | N/A | N/A | 476 |
|  | FPS | 43 | 53 | 81 | 104 | 125 | 148 | 172 |  |  | 0 |
|  | PPV% | 83.3 | 82.5 | 77.6 | 74.3 | 71.5 | 68.8 | 65.9 |  |  | 100 |
|  | TPR% | 45.0 | 52.5 | 58.8 | 63.2 | 66.0 | 68.7 | 70.0 |  |  | 100 |
| Low-Dose chromosome data | TPs | 13 | 18 | 18 | 20 | 20 | 20 | 20 | N/A | N/A | 27 |
|  | FPS | 37 | 51 | 67 | 90 | 120 | 136 | 156 |  |  | 0 |
|  | PPV% | 26.0 | 26.1 | 21.2 | 18.2 | 14.3 | 12.8 | 11.4 |  |  | 100 |
|  | TPR% | 48.2 | 66.7 | 66.7 | 74.1 | 74.1 | 74.1 | 74.1 |  |  | 100 |

<sup>1</sup> Results scored by University of Western Ontario (UWO/JHMK). DCs scored by UWO are treated as ground truth. Calculation of TPs and FPS based on comparing scoring by SVMs by HC and CNL are based on UWO ground truth. <sup>2</sup> The DC chromosome subset commonly scored by UWO, HC, and CNL and by the software was exposed to high dose radiation. <sup>3</sup> All data in the high-dose subset, scored by UWO and the software. This includes images that were not scored by all three experts. N/A: not applicable; TPs: true positive DCs; FPS: false positive DCs; PPV: positive predictive value; TPR: true positive rate

## Discussion

The overall accuracy of the DC detection algorithm relies on the combined performance of its three components: chromosome segmentation, centromere candidate assignment, and discrimination of DCs and MCs. However, image segmentation of metaphase chromosomes is not a trivial task. Under-segmentation hindered the performance of early releases of ADCI. Originally, the average number of segmented chromosomes (DC or MC) per image in dataset 1 was 12.4 (3488/281) and 24 (14429/531) in dataset 2. Both values are below the 46 chromosomes expected in a normal cell. Although inseparable chromosome clusters are eliminated by the software, reducing the TP DCs, this was preferable to the increased FP rates that would result from including these objects. Overlapping normal chromosomes (50%) are misclassified as DCs by commercial DCScore software (Metasystems; Vaurijoux et al. 2009) due to the presence of multiple centromeres per object. Application of the modified watershed transform largely resolved this problem for touching chromosomes or close neighbors (but not overlapping chromosome clusters). The watershed separation increased the average number of segmented objects per cell to near euploid levels, i.e. 38-40 per image (dataset 3). Although the modified Watershed algorithm handles homologous metaphases chromosomes with fused sister chromatids, it does promote over-segmentation in metaphase cells with severe sister chromatid separation or significant amounts of staining debris. Gaps between sister chromatids along the length of the chromosome create separate objects with variable intensity patterns resembling multiple discrete chromosome objects, which misleads watershed transform to produce ridges. Heuristically-designed conditional filters have been implemented to prevent over-segmentation (see Methods). Furthermore, the software avoids misclassification by selecting metaphase images by thresholding object counts per image. Excessive sister chromatid separation produces large numbers of segmented objects (>60) corresponding to individual chromatid arms rather than whole chromosomes. Using these object count thresholds, cells prone to DC misclassification due to over-segmentation can be eliminated.

The centromere detection algorithm has been optimized to reject false-negative DCs at the expense of higher false-positive rates. The method works well for identifying the first centromere (92% accuracy); however, detection of the second centromere based on the two highest ranked candidates is less accurate (70%). The candidates ranked and selected by the centromere SVM are important for making DC assignments. Incorrect centromere candidates affect the correct identification of true DCs by the MC-DC SVM. The current approach is approximately 70% accurate using the optimum σ values. Acrocentric chromosomes with short arms at the end of the DC or two acrocentric chromosomes forming DCs by fusion of their short arms are often misclassified as MCs (FNs). Centromere misclassification along chromatids is also common in SCS chromosomes. However, selecting centromeres among the 4 top-ranked candidates increases dicentric catchment rates. However, the preferred approach to train the MC-DC SVM with 4 centromere candidates has not yet been established.

One of the challenges in developing the centromere and MC-DC SVMs has been to develop image features that discriminated correct centromeres and DCs, independent of chromosome morphology. The most useful features were inspired by visual constrictions at centromeric structures and the corresponding width profiles. Other feature classes (F4 and F10) aimed at preventing or reducing FP DCs were discovered through review of testing results. A number of potential features in this class were ultimately not incorporated because of their minimal contribution or even adverse effect on accuracy. Some features are loosely defined, because of a lack of strict mathematical definitions for these biological characteristics. Examples include the curvature angles in F11-F15. The indexed distance of the 5-point offset to the Laplacian point on the contour used in the angle calculation was determined empirically, and validated to improve the accuracy of the MC-DC SVM through experimentation. We found that flexibility in these calculations has little effect on final classification results, as long as the results are biologically sensible. For instance, the steepness comparison of a pair of candidate width profiles, F9, which is measured by a relative ratio, can alternatively, be expressed as the absolute difference between these values without affecting the performance of the SVM.

The preferred SVM tuning parameters, σ, were empirically determined. There is a tradeoff between tuning the SVM to maximize either TPR or PPV (but not both). Increasing σ improves sensitivity, ie. more positive predictions of DCs, but reduces specificity. However, the numbers of MCs will always exceed DCs, regardless of radiation exposure. For this reason, the SVMs have been optimized to maximize correct detection of TP. σ values from 1.4 to 1.6 result in a balance of TP and FPs and maximize PPV and TPR. At high doses, at least, these sigma values provide satisfactory accuracy for differentiating MCs from DCs, though manual review by experts is more accurate when scoring is consistent.

At low dose exposure (<1 Gy), the algorithm identifies fewer DCs as expected. The FPR is near constant across a range of exposure levels, however small errors in DC detection at low dose will inflate dose estimation. The FPs are comprised of monocentric chromosomes, noisy objects and chromosome clusters or fragments that were not eliminated. Since there are multiple sources of FPs, no single solution may resolve this issue. One promising approach to reduce FPs involves normalization of image segmentation features of all chromosomes in a metaphase cell and using thresholding to discriminate outlier FPs relative to these normalized distributions.

To perform dose assessment will require constructing calibration curves from automated analysis of all DCs in a set of metaphase cells, and using these curves to predict doses for test samples processed using the same algorithms. Dose assessment comparisons between cytogenetic experts and the software will also be critical for adoption of automated approaches.

## Acknowledgements

We acknowledge support from the University of Western Ontario, Canada Research Chairs Secretariat, and the Canadian Foundation for Innovation. PKR and JHMK are founders of Cytognomix Inc., which has developed products for cytogenetics, and are inventors of US Patent No. 8,605,981

